# ATPIF1 inactivation promotes antitumor immunity through metabolic reprogramming of CD8^+^ T cells

**DOI:** 10.1101/2020.09.23.310979

**Authors:** Genshen Zhong, Ying Wang, Jiaojiao Zhang, Yichun Wang, Yuan Li, Yaya Guan, Shuang Shen, Xiaoying Zhang, Xinyu Cao, Minna Wu, Zhongxin Zhang, Ming Shi, Yunwei Lou, Yinming Liang, Hui Wang, Jianping Ye

## Abstract

Induction of CD8^+^ T cell activity is a promising strategy in the cancer immunotherapy. In this study, we identified ATP synthase inhibitory factor 1 (ATPIF1) as a potential target in the induction of CD8^+^ T cell immunity against tumor. Inactivation of ATPIF1 gene in mice promoted the antitumor activity of CD8^+^ T cells leading to suppression of tumor growth of B16 melanoma and Lewis lung cancer. The phenotype was abolished by deletion of CD8^+^ T cells in the ATPIF1-KO mice. The tumor infiltrating CD8^+^ T cells exhibited strong activities in the proliferation, effector and memory as revealed by the single cell RNA sequencing results of CD45^+^ tumor infiltrating lymphocytes (TILs) isolated from the tumors. The CD8^+^ T cells expressed more antitumor makers in the tumor microenvironment and in coculture with the tumor cells. The cells had a higher level of glycolysis after the T cell receptor-mediated activation as revealed by the targeted metabolomics assay. The cells exhibited an extra activity of oxidative phosphorylation before the activation as indicated by the oxygen consumption rate. The cells gained capacities in the proliferation, apoptosis resistance and mitophagy in the glucose-limiting environment. These data suggest that inhibition of ATPIF1 activity by gene inactivation rewired the energy metabolism of CD8^+^ T cells to enhance their immune activities to the tumors. ATPIF1 is a potential molecular target in the induction of antitumor immunity through metabolic reprogramming of CD8^+^ T cells for the cancer immunotherapy.

## Introduction

The application of PD-1/PD-L1 antibody revives cancer immunotherapy for advanced or drug-resistant cancers in the clinic practice. The success shed light on the strategy for metabolic reprogramming in the induction of immune activities of T cells ^1^, such as durability, longevity, and functionality ^2^. Energy supply is a primary factor in the control of those activities of T cells, especially in the glucose-limiting condition of the tumor microenvironment, in which T cells suffer energy deficiency due to lack of glucose and other nutrients ^3^. Glucose provides energy (ATP) and building materials to the T cells in support of cell survival and function ^4^. The energy shortage is intensified by the exhaustion proteins (PD-L1) of tumors ^2^. Metabolic reprogramming is a potential strategy to overcome the energy shortage in T cells by maintaining the energy homeostasis ^5^. However, the strategy is limited by a lack of the molecular targets in the metabolic reprogramming ^4, 6^.

ATPIF1 is an inhibitory protein of F_1_F_o_-ATP synthase (ATP synthase, Complex V) in mitochondria, which catalyzes ATP production through phosphorylation of ADP at expense of mitochondrial potential in the energy substrate-enriched environment. The ATP synthase hydrolyzes ATP in the energy substrate-deficient conditions to protect cells from apoptosis by maintaining the potential. Both activities of ATP synthase are regulated by ATPIF1 ^7, 8^, which involves a protein-protein interaction ^9^. ATPIF1 is highly expressed in mitochondrion-enriched cells to match the ATP synthase activity. An increase in ATPIF1 activity in the ATPIF1 overexpression mice, the ATP synthase activity is decreased leading to energy metabolism reprogramming for down-regulation of oxidative phosphorylation (OXPHOS) and up-regulation of glycolysis in mice ^7^. The reprogram increases the risk of cell apoptosis under conditions of respiration inhibition and energy deficiency from the mitochondrial potential collapse ^10^. ATPIF1 activity has been investigated in the regulation of metabolism in the neuronal cells ^11^, red blood cells ^12^, epithelial cells ^13^, hepatocytes ^14^ and tumor cells ^15^. However, its role remains unclear in lymphocytes.

In this study, we observed that ATPIF1 inactivation promoted the CD8^+^ T cells activities in proliferation, survival and effector memory as indicated by single cell RNA sequencing assay of CD45^+^ leukocytes isolated from the B16 tumor. The impact was a consequence of metabolic reprogramming in CD8^+^ T cells following an elevation of the ATP synthase activity. These results suggest that APTIF1 is a potential molecular target in the induction of anti-tumor activity of CD8^+^ T cells by metabolic reprogramming.

## Results

### Antitumor immunity is enhanced in ATPIF1-KO mice in dependence on CD8^+^ T cells

The ATPIF1-KO (ATPIF1^-/-^) mice was generated using the CRISPR-Cas9 strategy, and the inactivation was verified with a decrease in ATPIF1 protein in the spleen lymphocytes by Western blotting (Fig. 1a). The knockout mice exhibited no difference from the wild type (WT) mice in growth and reproduction. The impact of ATPIF1 inactivation in the immune function remained unknown. We addressed this issue by analyzing the percentage of T cells, B cells, Treg cells, dendritic cells, and neutrophil cells in the cell population of ATPIF1-KO mice spleen. No significant alteration was detected in those cells in the normal condition (Fig. S1), but the percentage of CD11c^+^ dendritic cells was increased in the population analysis (Fig. S1c).

**Figure 1.**
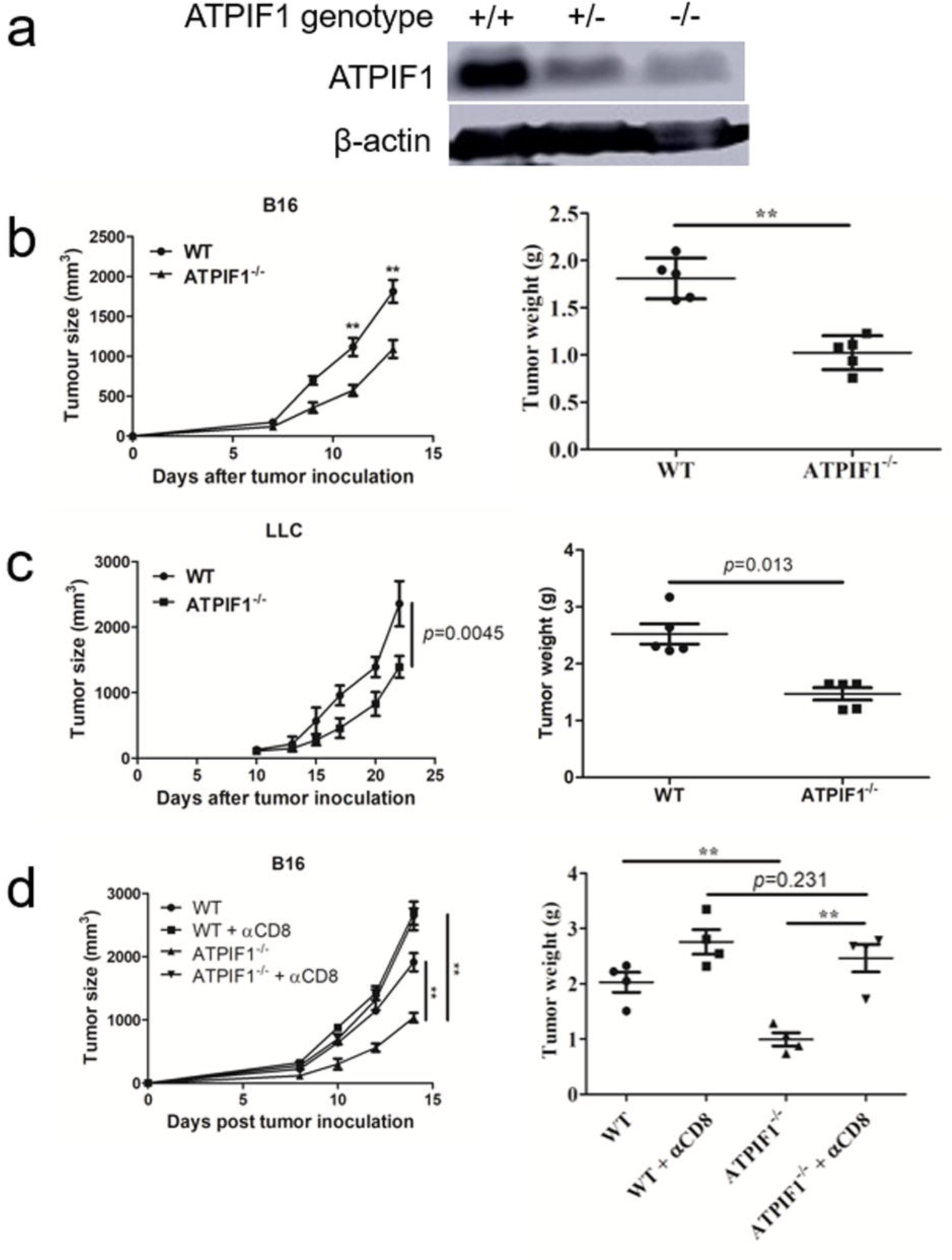
Antitumor activity is enhanced in ATPIF1^-/-^ mice with dependence on CD8^+^ T cells. (a) Confirmation of ATPIF1 deletion in the KO mice. ATPIF1 protein was determined in the mouse spleen by WB to confirm the gene inactivation in lymphocytes. (b) Reduction of B16 melanoma cancer growth in ATPIF1^-/-^ mice. (c) Reduction of Lewis lung cancer growth in ATPIF1^-/-^ mice. The tumors were engrafted subcutaneously and examined for tumor weight on day 14 and 24, respectively. (d) Dependence of tumor immunity on CD8^+^ T cells. CD8^+^ T cells were deleted in the mice with intraperitoneal injection of the CD8 antibody (200 μg/per mouse) twice/week during the B16 melanoma tumor growth period. Tumor growth was determined at day 14 after engraftment. The tumor weight was presented as mean ± SD. ** *p*<0.01.

To investigate the immune system further, we examined the antitumor activity of KO mice in the two tumor-bearing models of B16 melanoma and Lewis lung cancer (LLC). The B16 tumor growth was significantly reduced after engraftment into the KO mice with a smaller tumor size at two weeks. A 56.6% reduction in the tumor weight was observed (*p* = 0.0017, Fig. 1b). A similar reduction in tumor size was found in LLC tumor in the KO mice for 41% decrease in tumor wight relatively to that of WT mice (*p* = 0.0045, Fig.1c). To elucidate the T cell activity in the phenotype, the CD8^+^ T cells were depleted in the mice using a CD8 antibody during the B16 tumor growth (Fig. S2). The depletion led to a robust acceleration in tumor growth in the WT and KO mice (Fig. 1d), which abolished the difference between the two groups of mice in tumor growth. These data suggest that ATPIF1 inactivation enhanced the tumor immunity in the KO mice to inhibit growth of engrafted tumors, which was dependent on CD8^+^ T cells.

### ATPIF1 inactivation favors CD8^+^ T cell expansion in TILs

Above data suggest that the ATPIF1 inactivation may generate an impact in CD8^+^ T cells to enhance the tumor immunity. To test the possibility, the single-cell RNA sequencing (scRNA-seq) technology was used in the analysis of TILs heterogenicity. The scRNA-seq technology has been reported in the landscape analysis of TILs of hepatocellular carcinoma ^16^, breast cancer ^17^ and colorectal cancer ^18^. The assay was conducted using CD45^+^ cells isolated from the tumor tissues with 8408 cells from the KO mice and 10059 cells from the WT mice (Fig. S3). The sequencing data were obtained from 6007 leukocytes in the KO group and 8010 leukocytes in the WT group based on the 10x genomics method (Table S1). The violin chart and the principal component analysis (PCA) indicated the well quality control during the scRNA-seq (Fig. S4a and S4b), also, the T-distributed random neighborhood embedding (tSNE) and uniform manifold approximation and projection (UMAP) graph showed the similar landscape between the WT and KO sample (Fig.S4c and S4d).

ATPIF1 inactivation promoted maturation of naïve T cells and proliferation of T cells to increase the percentage of CD8^+^ T effector (T_eff_) and T memory (T_mem_) cells in TILs. Among the CD45^+^ leukocytes, 1353 CD3^+^ T cells in the KO sample and 907 CD3^+^ T cells in the WT sample were finally sequenced. Eight clusters of CD3^+^ T cells were identified in TILs by scRNA-seq (Fig. 2a) and the marker genes in each cluster was summarized (Fig.S5). In the KO mice, the clusters of naïve T cells (KO:WT=39.4%:57.8%) and CD4^+^ T_reg_ (KO:WT=5.54% 7.72%) were decreased. The clusters of proliferative T cells (KO:WT=9.02%:3.31%), CD4^+^ T cells (KO:WT= 21.7%:12.4%) and CD8^+^ T_eff_ and T_mem_ cells (KO:WT=8.80%:3.64%) were increased (Fig. 2b). The proliferation was mainly found in the CD8^+^ T cells by the markers of MKI67 and Stmn1 (Fig. 2c). IFN-γ mRNA was increased in T cells of KO mice (Fig. 2d). These indicate an enhancement in the proliferation and effector (IFN-γ secretion) capacities in T cells of the KO mice. The STARTRAC (single T cell analysis by RNA-seq and TCR tracking) program ^18^ was employed to track the dynamic relationships among T cell subsets in TILs. STARTRAC is a novel analytical framework in investigation of the clonal expansion, migration and state transition of TILs, which provide critical insights into the T cell lineages ^18, 19^. In the KO mice, the CD8^+^ T_eff_ and T_mem_ cells exhibited an enhanced clonal expansion, which was followed by expansion of T-proliferative cells (Fig. 2, e and f). In contrast, the expansion of CD4^+^ Treg was predominant in the WT mice (Fig. 2g), which was associated with expansion of CD4^+^ T cells (Fig. 2h). CD4^+^ Treg mediates the immunosuppression activity in tumor ^6^. The pseudotime trajectory analysis was used to characterize the lineage differentiation of TILs. The developmental trajectory from the T proliferative cells to CD8^+^ T_eff_ was enhanced in the KO mice (Fig. 2i). To verify the result, we analyzed the CD8^+^ T_eff_ and T_mem_ in the peripheral blood mononuclear cells (PBMC) and TILs using flowcytometry. The percentage of T_eff_ (CD44^+^ CD62L^-^ CD8^+^) and T_mem_ (CD44^+^ CD62L^+^ CD8^+^) was increased dramatically in the KO mice, especially in TILs (Fig. 2j).

**Figure 2.**
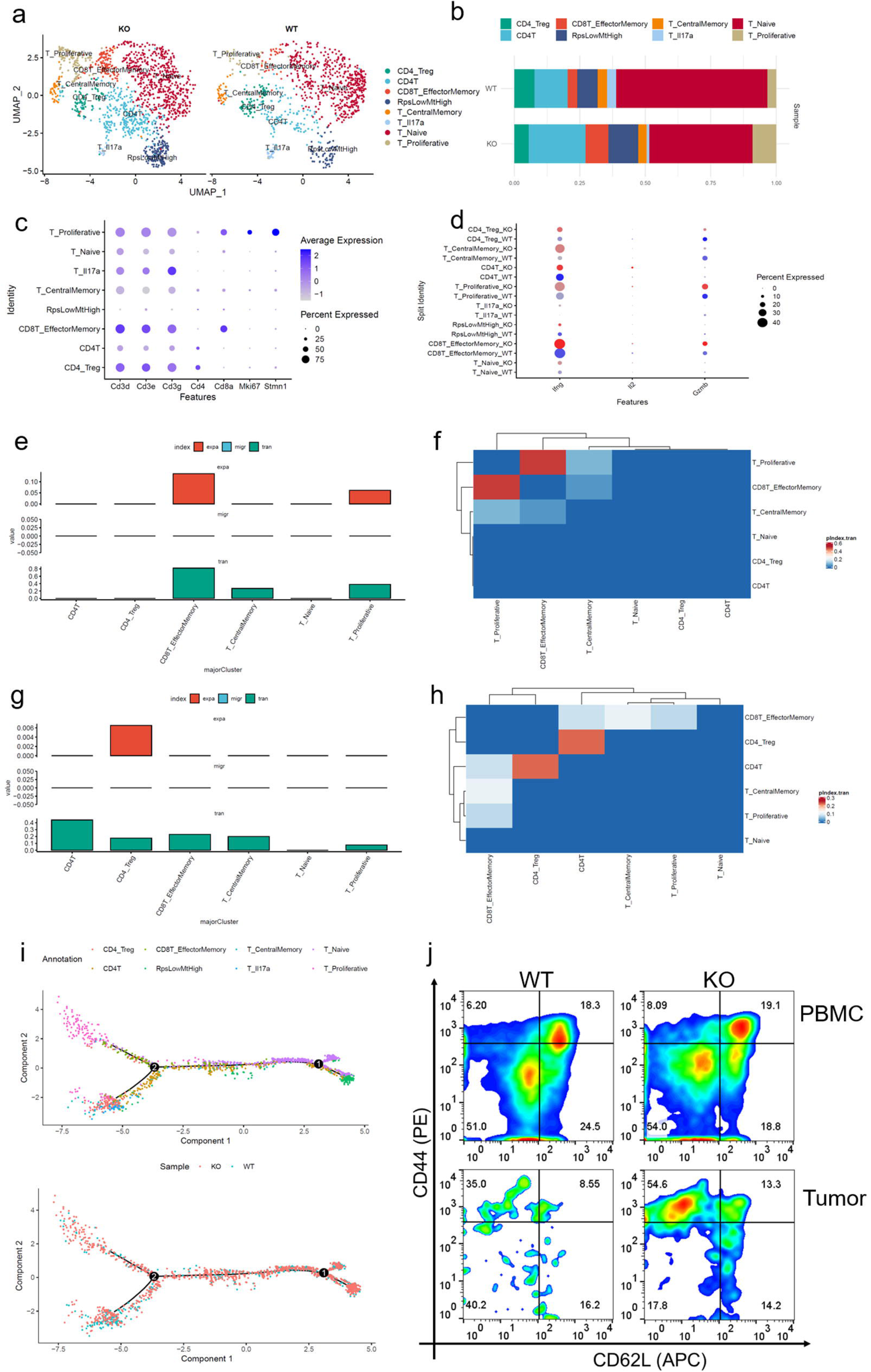
ATPIF1-deficiency enhanced the CD8^+^ T cell proliferation and the formation of memory CD8^+^ T cell in tumor environment. (a) t-SNE cluster map of T cells based on the canonical markers, eight subclusters were graphed. (b) Percentage of different clusters of T cells in the tumors of WT and KO mice. (c) Strongest proliferation capacity of CD8^+^ T cells based on the analysis of Mki67 and Stmn1. (d) Increased IFN-γ mRNA content in T cells of KO over WT mice. (e) and (f) Strong expansion and transition potential in CD8^+^ Teff/Tmem based on the TCR sequencing using the STARTRAC analysis approach, which reflects the T cell expansion (exp), migration (migr) and transition (tran). (g) and (h) The high expansion potential of Treg cells in WT mice and closely related with CD4^+^ T cells based on the STARTRAC method. (i) The pseudotime trajectory analysis of T cells based on integrated expression and TCR clonality. The order of T cells along pseudotime is expressed in a two-dimension state-space defined by Monocle2. Cell orders are inferred from the expression of most dispersed genes across T subsets sans MAIT. Each point indicates a single cell, and each color represents a T cell cluster. (j) Flow cytometry analysis of CD8^+^ memory T cells in PBMC and tumor single cell suspension, which were collected from the B16 tumor of WT and KO mice. CD8^+^ Teff/Tmem were defined as CD44^+^CD62L^-^ CD8^+^ T cells.

### Function of CD8^+^ T cells is enhanced in KO mice under the glucose-limiting condition

Above data suggest that the CD8^+^ T cells exhibited an enhanced proliferative capability and effector memory activity in the KO mice. To verify the results, CD8^+^ T cells were investigated in proliferation and IFN-γ secretion in the cell culture in vitro. The cells were isolated and activated with T cell receptor signal mimics (CD3 plus CD28) in the culture. The test was conducted with three concentrations of glucose as glucose is required for T cell function and IFN-γ production ^20^. Interestingly, the KO cells had a delayed proliferation under the high (HG) and normal (NG) levels of glucose (Fig. 3a). However, the phenomenon was reversed under the low glucose (LG) condition, which resembles the glucose-limiting environment in tumor. Consistently, IFN-γ secretion was increased in the KO cells under the low glucose condition (Fig. 3b). Glucose uptake was examined in T cells using a fluorescent glucose analog (2-NBDG) to access the glucose metabolism ^21^. In the KO cells, the uptake was significantly increased in the LG condition (Fig. 3, c and d), which was observed with more mRNA expression of glucose transporter 1 (Glut1, slc2a1) (Fig. 3e). mRNA of pyruvate dehydrogenase kinase 1 (PDK1) was upregulated in the KO cells in the same condition (Fig. 3f), suggesting an inhibition of the oxygen-dependent glucose metabolism. The data suggest that T cell proliferation, IFN-γ production and glucose uptake were all enhanced in the KO cells under the low glucose condition.

**Figure 3.**
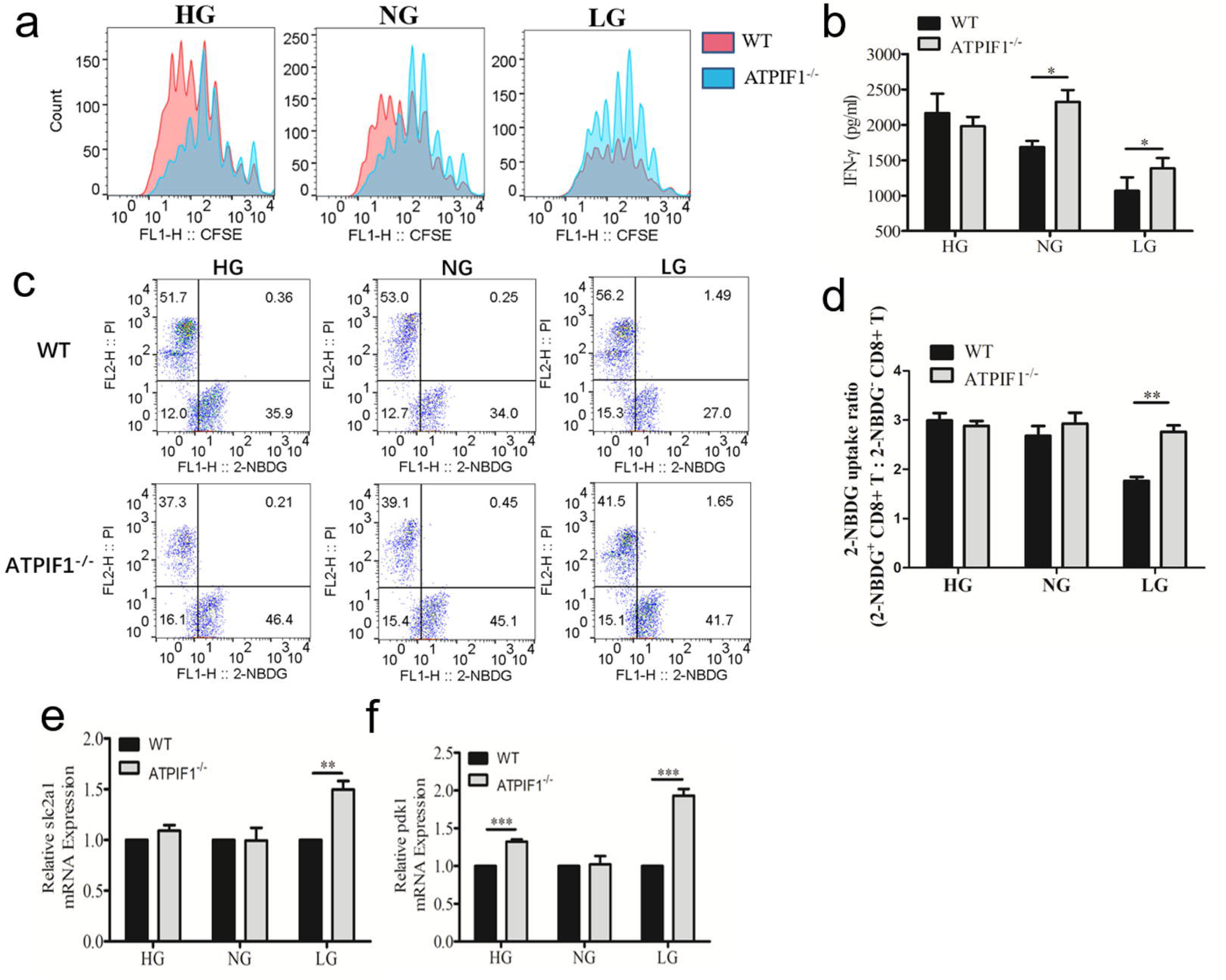
Function of CD8^+^ T-cells of KO mice is enhanced in the low glucose condition. (a) CD3^+^ T-cells were isolated from the spleen and labeled with CFSE. The cells were stimulated with CD3/CD28 dynabeads in the culture medium with different glucose concentrations (HG, NG, and LG). Analysis was conducted 4 days later with staining by CD3/CD8 antibodies. Light-red: WT and Light-blue: ATPIF1^-/-^. (b) IFN-γ in the supernatant determined with ELISA. (c) Glucose uptake. T-cells were cultured under HG, NG, and LG conditions for 24 h, then examined for glucose uptake with 2-NBDG treatment for 1 h at 37□. Staining for CD3/CD8 and Propidium Iodide (PI) was conducted for the CD8^+^ T cells. (d) Statistical analysis of 2-NBDG positive cells. The 2-NBDG positive cells were calculated in the CD8^+^ population. (e) Glut 1 protein. The protein was determined by Western blotting in CD8+ T-cells following culture in the HG, NG, and LG conditions for 24 h. (f) Glut 1 mRNA. The Glut 1 encoding gene slc2a1 was examined in mRNA with qRT-PCR in CD8+ T-cells, which were cultured in the HG, NG, and LG conditions for 24 h. (g) Pdk1 mRNA. The mRNA was determined in the same condition above. Data are presented in mean ± SD (n=3), *p<0.05, **p <0.01, ***p<0.001.

### Glycolysis is enhanced in KO cells upon activation by CD3 plus CD28 signals

Glucose is the major fuel in T cells in the production of ATP, which involves two steps: glycolysis and oxidative phosphorylation (OXPHOS). The two steps were examined in CD8^+^ T cells using the Seahorse equipment for the oxygen consumption rate (OCR) of OXPHOS, and the extracellular acidification rate (ECAR) for aerobic glycolysis, respectively. In the KO CD8^+^ T cells, OXPHOS was more active in the resting state (or naive state) (Fig. 4a, *p*<0.05), but less active in the activated state (Fig. 4b, *p*<0.05). The spare respiratory capacity (SRC) was increased in both resting and activated conditions (Fig. 4, a and b). Glycolysis was less active in the resting state as indicated by the low value of ECAR (Fig. 4c, *p*<0.05), which became more active after cell activation for the rise in ECAR value (Fig. 4d). The glycolytic reserve (GR) was significant higher in the KO cells in both resting and activated states (Fig. 4, c and d, *p*<0.01). The increase in SRC is a character of energy metabolism in the memory antitumor T cells, and GR increase is a character of the effector antitumor T cells ^31, 32^. The data suggest that glucose metabolism is reprogrammed in CD8^+^ T cells of KO mice in favor of an energy adaptation to the antitumor activity.

**Figure 4.**
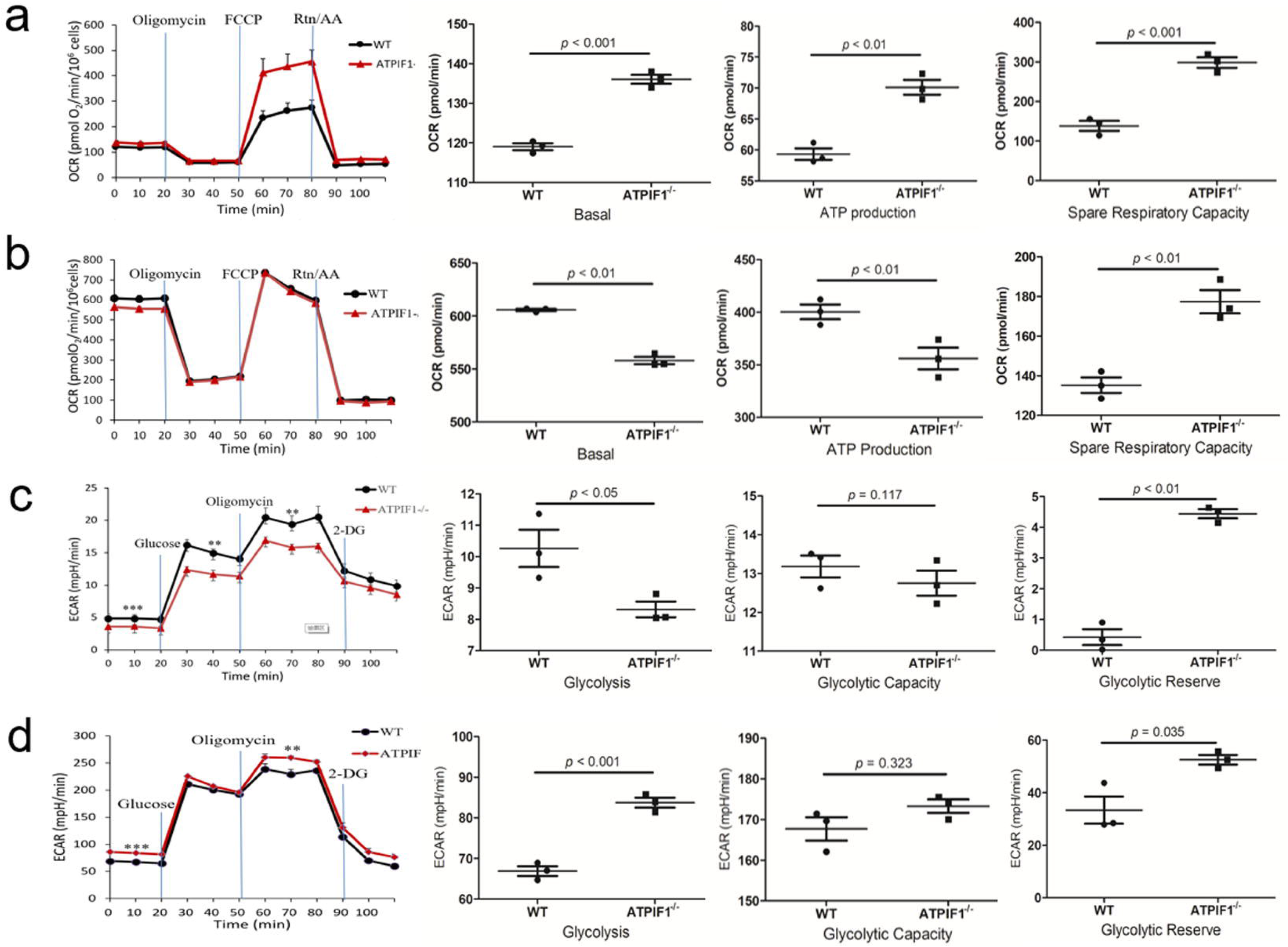
KO cells are reprogramed for active glycolysis upon activation by CD3 plus CD28 signals. (a) Oxygen consumption rate (OCR) in resting cells. CD8^+^ T cells were examined for mitochondrial activity with OCR at the baseline (without CD3/CD28 antibody stimulation). OCR in the activated cells. The test was conducted in CD8^+^ T-cells after CD3/CD28 co-stimulation for 24 h in the culture. ATP production and Spare Respiratory Capacity (SRC) of the cells in basal or activated states were calculated and plotted. (c) Extracellular acidification rate (ECAR) in resting T-cells. (d) ECAR in activated T-cells. The glycolysis, glycolytic capacity and glycolytic reserve were calculated at the basal or activated states of cells. Data represents mean ± SEM (n=3). * *p*<0.05, ** *p*<0.01, *** *p*<0.001.

### Targeted metabolomics reveals an elevation in glycolysis and pentose phosphate pathway in the KO T cells

Targeted metabolomics was used to investigate the metabolic reprogramming in the KO cells. CD8^+^ T cells (10^7^) were purified from the spleen of tumor-bearing (B16-OVA) mice through FACS-mediated sorting. Metabolites in the energy metabolism pathways were analyzed using the liquid chromatography-mass spectroscopy (LC-MS). The metabolites include those in the tricarboxylic acid cycle (TCA), glycolytic pathway, OXPHOS and pentose phosphate pathway. The metabolite profile revealed a higher activity in the glycolysis and the pentose phosphate pathways for the increase of D-Glucose 1-phosphate, D-Glucose 6-phosphate, α-D-Ribose 5-phosphate and coenzyme A (Fig. 5, a and b, raw data are summarized in Table S2). The TCA cycle activity was reduced for the decrease in acetoacetyl-CoA and acetyl-CoA. The changes in metabolite profiles favor the production of antitumor cytokines, such as IFN-γ in T cells ^29^. The data suggest that CD8^+^ T cells of KO mice were more active in adaptation to aerobic glycolysis from OXPHOS in the fight against tumor.

**Figure 5.**
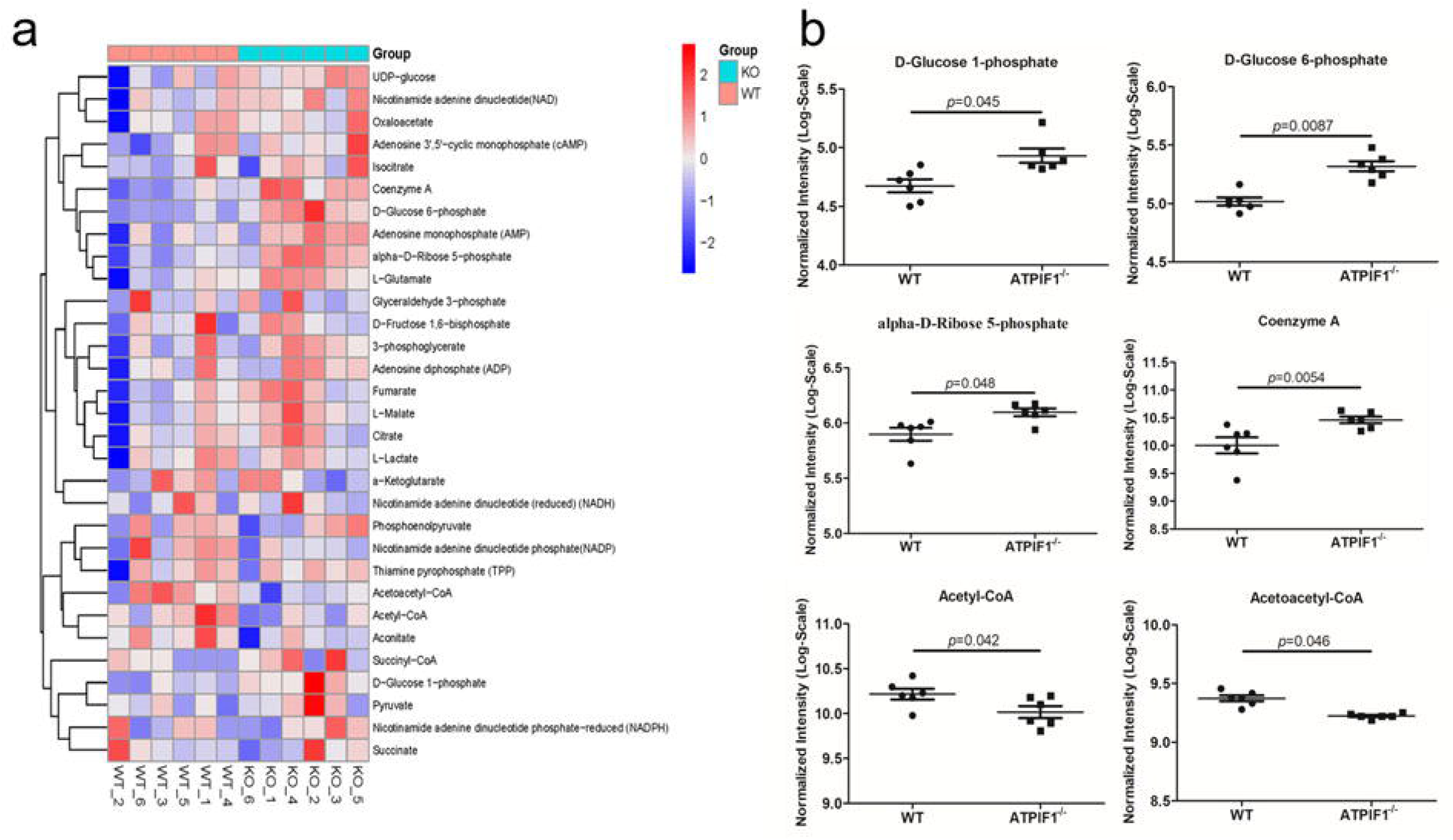
Targeted metabolomics reveals an increase in metabolites of glycolysis and pentose phosphate pathway in the KO cells. (a) Heatmap of targeted metabolomics focus on the 31 intermediate products of glucose in the tricarboxylic acid cycle (TCA), glycolytic pathway, OXPHOS and pentose phosphate pathway in CD8^+^ T cells of KO vs. WT mice. The CD8+ T cells were isolated from the spleen of B16-OVA tumor-bearing mice and analyzed as described in the Methods section (n=6). (b) Quantification of 6 intermediate products in the targeted metabolomics. The data represents mean ± SD (n=6), * *p*<0.05, ** *p*<0.01.

### CD8^+^ T cells exhibit an enhanced capacity in the maintenance of mitochondrial potential (ΔΨm) in the KO mice

The F_1_F_o_-ATP synthase synthesizes as well as hydrolyzes ATP, which will rise in the absence of ATPIF1 according to studies in other cell types. However, the activity alteration remains unknown in T cells of KO mice. To address the issue, the CD8^+^ T cells were examined with several parameters including the intracellular ATP level, mitochondrial potential (ΔΨm), ROS, and MitoSOX. The tests were conducted in the cells under different glucose concentrations to understand the alteration in ATP synthase activities. The intracellular ATP content was increased in the KO CD8^+^ T cells at the high (HG) and normal (NG) glucose concentrations to support an increase in the synthetic activity of ATP synthase in the energy sufficient conditions (Fig. 6a). The ATP was decreased at the low glucose (LG) concentration to reflect the rise in hydrolytic activity of ATP synthase in the energy deficient condition (Fig. 6a). The hydrolytic activity is required for the maintenance of ΔΨm under the energy deficient condition ^33^. ΔΨm was examined in the cells using JC-1 and the level was higher in the KO cells over the WT cells at the LG condition (Fig. 6b). The difference was also observed in the HG condition. ΔΨm was decreased in the LG condition in both WT and KO cells, but the reduction was attenuated in the KO cells. The data suggest that the hydrolytic activity of ATP synthase was increased more efficiently in the KO cells under the energy deficient condition to consume ATP in the maintenance of ΔΨm.

**Figure 6.**
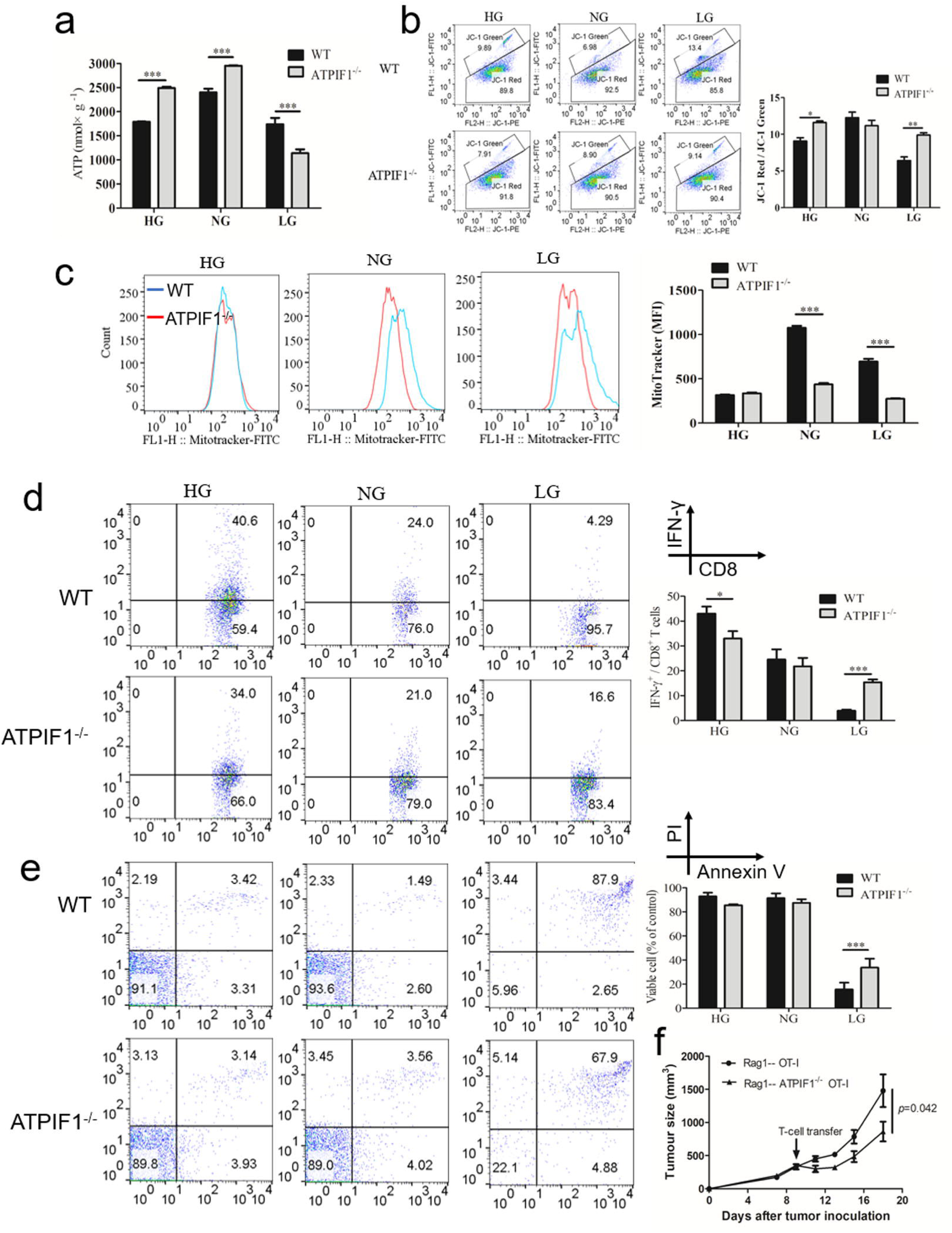
KO cells exhibits an enhanced energy metabolism capacity in the low glucose condition. (a) Intracellular ATP content. CD8^+^ T cells were cultured under the HG, NG, and LG conditions for 24 h, and then examined for the ATP content using the luciferin-based assay. (b) Mitochondrial membrane potential. The ratio of JC-1 red to JC-1 green was calculated for the potential change. (c) Mitochondrial mass. The signals were quantified with flow cytometer and the mean fluorescence intensity (MFI) was calculated and plotted on the right bar figures. CD8^+^ T cells were cocultured with B16-OVA tumor cells in a ratio of 1:1 under different glucose concentrations. Analysis was conducted in the T-cells after 36 h using the flow cytometry. (d) IFN-γ content. (e) Apoptosis assay. (f) Transfer of ATPIF1^-/-^ CD8^+^ T cells to the B16-OVA xenografted Rag1 mice. Rag1 mice were inoculated with 2×10^6^ B16-OVA melanoma cells to establish the tumor-baring model, 9 days later, 2×10^6^ of ATPIF1^-/-^ OT-I or OT-I CD8^+^ T cells were intratumor injection to test the antitumor immunity. Nine days later, the mice were sacrificed for tumor size analysis (n=5). Data represents mean ± SD. *p<0.05, **p<0.01, ***p<0.001.

The mitochondrial mass was determined using MitoTracker staining. The mass was decreased in the KO CD8^+^ T cells in the normal (NG) and low (LG) glucose conditions (Fig. 6c). These data demonstrate that the ATP synthase activity was reprogrammed in CD8^+^ T cells of ATPIF1-KO mice for an improved adaptation to the energy supply.

### Immune activity of CD8^+^ T cells is enhanced in KO mice in vitro and in vivo

The antitumor function of CD8^+^ T cells was tested further for IFN-γ expression *in vitro* and *in vivo*. The study was conducted with OT-I mice, in which the CD8^+^ T cells are engineered to express a T cell receptor (TCR) motif for recognition of ovalbumin antigen of SIINFEKL peptide ^34^. ATPIF1 was inactivated in the OT-I mice by crossing the KO mice with OT-I mice. The CD8^+^ T cells was isolated from the hydride mice (ATPIF1^-/-^ OT-I) and challenged with the ovalbumin-expressing tumor (B16-OVA) cells to induce IFN-γ expression *in vitro*. Intracellular IFN-γ was quantified in the T cells using flow cytometry. IFN-γ was elevated in the cells by HG and decreased in the cells by LG in the WT mice (Fig. 6d). However, the LG response were significantly attenuated in the KO cells (Fig. 6d) for much more IFN-γ production in the low glucose condition. The KO cells were resistant to apoptosis under the LG condition (Fig. 6e, *p*<0.001). The data suggest that CD8^+^ T cells of ATPIF1-KO mice exhibited a much better activity in the INF-γ production and apoptosis resistance under the glucose-limiting condition. To confirm the result *in vivo*, the CD8^+^ T were isolated from the OT-1 mice or ATPIF1^-/-^ OT-1 mice using the Dynabeads™ of CD8^+^ positive isolation kit and then transferred into the B16-OVA xenografted Rag1 mice via the intratumor injection. The tumor inhibition was significantly improved in the ATPIF1^-/-^ group over the WT group (*p*=0.042) (Fig. 6f), supporting the enhanced antitumor efficacy of CD8^+^ T cells of ATPIF1-KO mice.

### Mitophagy is enhanced in CD8^+^ T cells of KO mice under the glucose-limiting condition

In the glucose-limiting condition, the survival ability was enhanced in the CD8^+^ T cells of ATPIF1-KO mice to support the immune function. These suggest that the KO cells may gain extra ability to obtain energy beyond glucose. Autophagy was examined to test the possibility by examining the representative markers. In the KO cells, mitophagy (PINK1 and Parkin) and autophagy (ATG5) markers were significantly increased (Fig. 7, a and b). The increases were observed under the LG as well as NG condition. The increases occurred regardless of activation status in the CD8^+^ T cells. Consistently, more autophagic vacuoles were found in the cytoplasm of KO cells under the electronic microscope (Fig. 7c). Mitophagy was induced in the normal (WT) T cells following activation by CD3+CD28 under the LG condition, and the response was further enhanced by the mitochondrial uncoupler CCCP (an mitophagy agonist) (Fig. 7d). Similar responses were observed in the KO cells in the presence of higher level of basal mitophagy (Fig. 7d). The enhanced autophagy was associated with more IFN-γ expression in the KO cells after activation by CD3/CD28 (Fig. 7e). However, the CCCF-induced IFN-γ response was attenuated in the KO cells (Fig. 7e). These data suggest that mitophagy was increased in the KO cells to provide energy in the glucose-limiting conditions.

**Figure 7.**
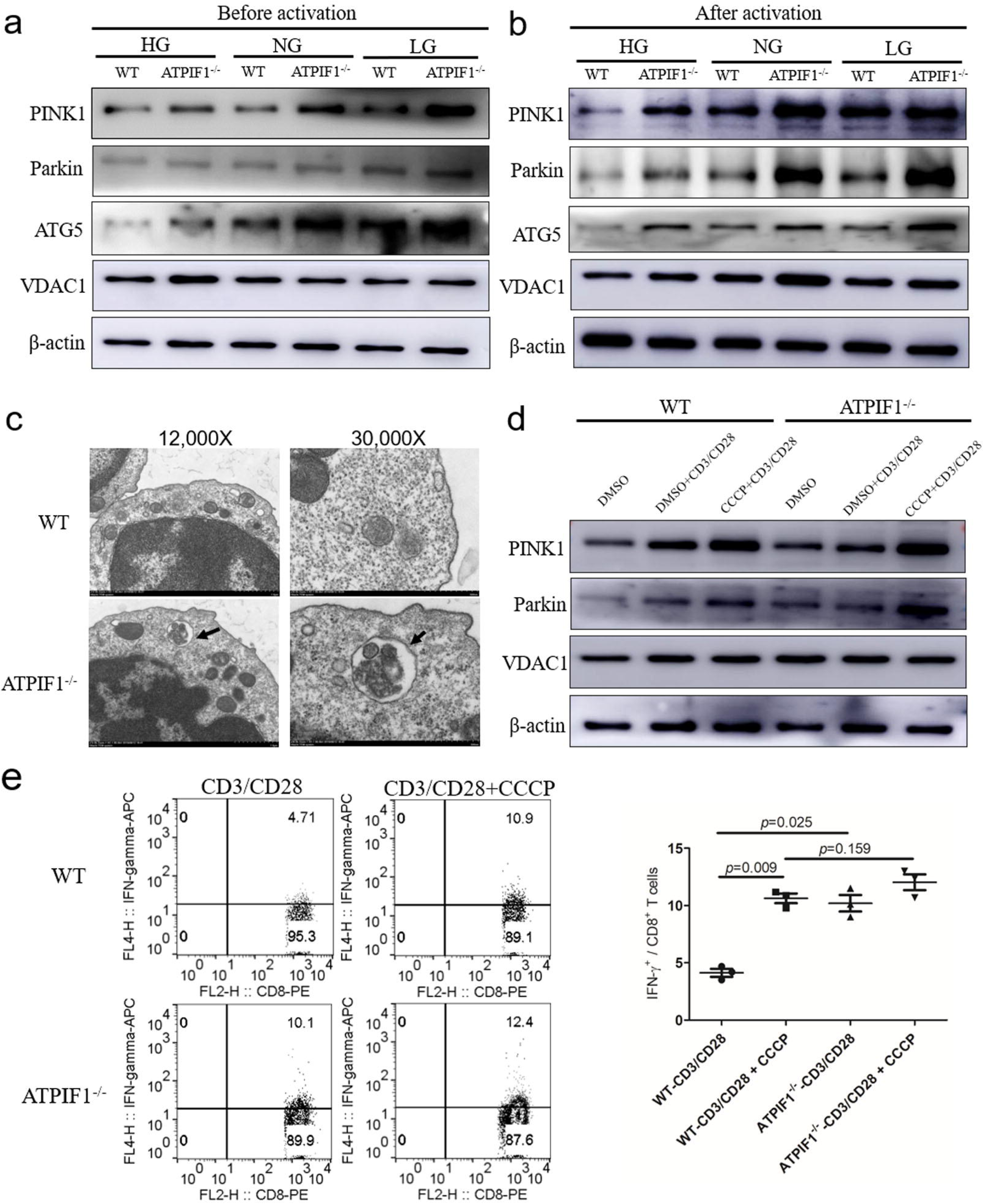
Mitophagy is enhanced in CD8^+^ T cells of KO mice under the glucose-deficient condition. Protein expression of PINK1, Parkin, ATG5, and VDAC1 were examined in CD8^+^ T cells under the HG, NG, and LG conditions. (a) Mitophagy in the resting T-cells. (b) Mitophagy in the activated T-cells by CD3/CD28 co-stimulation. (c) Electronic microscope image. Mitophagysome was observed in the cytoplasm of CD8^+^ T cells under the low and high magnification. Black arrows indicate the location of autophagic vacuole. (d) Mitophagy markers. The markers were examined in T cells treated with the uncoupler CCCP in the activated cells with CD3/CD28 stimulation under the LG condition. (e) Promotion of IFN-γ expression by mitophagy. The expression was induced in T-cells by CD3/CD28 co-stimulation. CCCP was used to induced mitophagy in the activated T-cells to promote the IFN-γ expression. The data represents mean ± SD (n=3). **p<0.01.

### Macrophage and dendritic cell clusters in TILs support the enhanced tumor immunity in KO mice

The alteration in macrophage and dendritic cell (DC) clusters of TILs supports the enhanced tumor immunity in the KO mice. In the scRNA-seq assay, four major clusters and 18 subclusters were identified in TILs based on the gene expression profiles (Fig. 8a). The major clusters included T cells, macrophages, B cells, and dendritic cells by the canonical marker genes (Fig. S6), indicating a well coverage of the cluster spectrum. Five macrophage clusters were identified in TILs (Fig. 8a). The macrophage_S100a8 cluster was decreased dramatically in the KO mice by 75% (Fig. 8b). Macrophage_S100a8 is an indicator of poor clinical outcome in the breast cancer for tumor invasion and migration ^22^. The immune suppression-related genes including Socs3, Plaur (urokinase plasminogen activator receptor, uPAR), and Arginase 2 (Arg2) were significant decreased in the cluster in the volcano plot (Fig. 8c), supporting that the immunosuppressive activities of macrophages were reduced in the KO mice ^23, 24, 25^. In the other four clusters, three of them such as macrophage_Lars2, macrophage_Apoe, and macrophage_Dct were increased, and one of them, Macrophage_CellCycle, was not altered in the KO mice (Fig. 8b). Functions of those four clusters are not known yet in the tumor immunity.

**Figure 8.**
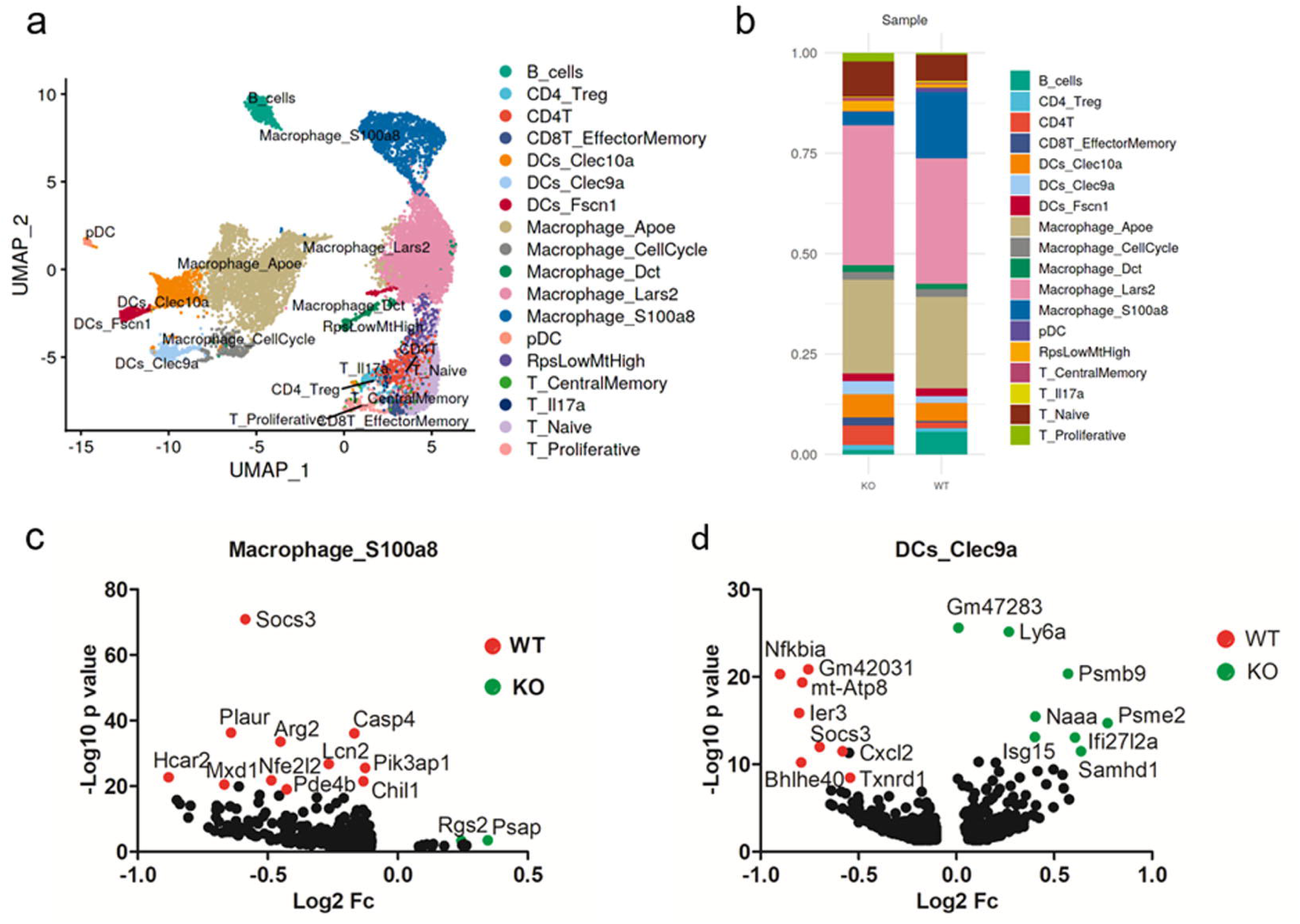
Macrophage and dendritic cell clusters support the improved antitumor immunity of TILs by scRNA-seq analysis. (a) Clustering of CD45^+^ leukocytes and distribution of selective functional-related genes expression. Four major clusters and 18 subclusters were identified in TILs. (b) Percentage of different clusters of TILs in the WT and KO samples. (c) and (d) Volcano plot of highly expressed genes in macrophage_S100a8 and DCs_Clec9a in the WT and KO samples, respectively. Red dots indicate genes that were highly expressed in the WT sample and green dots indicate genes that highly were expressed in the KO sample.

The enhanced immunity is further supported by the alteration of dendritic cells (DCs) (Fig. 8, a and b). Four clusters of DCs (Clec9a^+^ DCs, Clec10a^+^ DCs, plasmacytoid DCs and Fscn1^+^ DCs) were identified in TILs by the scRNA-seq assay. In the KO mice, there were 100% increase in Clec9a^+^ DCs (KO:WT=3.35%:1.72%), and 25% increase in Clec10a^+^ DCs (KO:WT=5.74%:4.50%). Differentiation of Clec9a^+^ DCs was enhanced in the potential developmental trajectory data (Fig. S7). Clec9a is an important marker of DCs in the recognition of necrotic cells and cross-presented antigens ^26, 27^. In this cluster DCs, the genes including Socs3 and IκBα (Nfkbia) were decreased significantly together with Gm42031, mt-Atp8, and Ler3 (Fig. 8d). Clec10a is an endocytic receptor for maturation and initiation of immune response in DCs ^28^. Plasmacytoid DCs (pDCs) produce type I interferons in the antiviral immunity, and its function is multifaced in the tumor immunity ^29^. A 2-fold decrease (KO:WT=0.33%:1.10%) was found in the cluster of pDCs. The DC cluster profiles further support the enhanced tumor immunity in TILs of the KO mice.

## Discussion

Current study suggests that ATPIF1 is a new candidate of molecular target in the induction of antitumor activity of CD8^+^ T cells. The antitumor activities of CD8^+^ T cells were enhanced in the ATPIF1-KO mice, which represents a novel observation. ATPIF1 activity has been studied in several types of cells, but not yet in lymphocytes. In the knockout studies, ATPIF1 inactivation enhanced pro-survival activity of hepatocytes by inhibition of apoptosis in the respiratory chain toxification models ^10^, and attenuated the pressure-induced cardiomyocyte hypertrophy ^30^. The inactivation led to an increase in ATP production by OXPHOS in macrophages ^31^ and promotion of adipocyte differentiation ^32^. In the overexpression studies, ATPIF1 activation reduces mitochondrial ATP production in the neuronal cells ^11^, intestinal epithelial cells ^13^ and hepatocytes ^14^. The activation leads to more ROS production by mitochondria for a high risk of pro-inflammatory response in the brain ^11^ and gut ^13^. The activation raises the glycolytic activity in favor of hepatoma growth ^14^. In the red blood cells, ATPIF1 is required for haem synthesis during the erythroblast differentiation in the hematopoietic models ^12^. However, an impact of ATPIF1-deficiency or activation in the immune system has not been reported in the literature. Current study provides evidence that ATPIF1 inactivation led to an increase in the tumor immunity in the tumor-baring mice. Deletion of CD8^+^ T cells abolished the antitumor immunity in the ATPIF1-KO mice. The data demonstrates a role of ATPIF1 in the regulation of CD8 T cells.

The enhanced activities of CD8^+^ T cells are supported by the cluster analysis of TILs with scRNA-seq and STARTRAC method, which reveal the subset landscape, the developmental trajectory, and TCRs clonal lineage in TILs. In this study, scRNA-seq was employed to determine the CD8^+^ T cell activity in TILs, in which eight major subsets were identified. In the KO mice, the T-proliferative and CD8^+^ T_eff_/T_mem_ clusters were increased, while the naïve T cell cluster was decreased. The CD8^+^ T clusters were most active in proliferation among the T cell subsets. To delineate the dynamic relationships of different T cell subsets, the STARTRAC indices were established in TILs to measure clone expansion. The CD8^+^ T_eff_ /T_mem_ cluster was most active in the clone expansion. These suggest that activities of CD8^+^ T cells were enhanced in the proliferation and differentiation of T_eff_/T_mem_ in the KO mice. These data support a role of ATPIF1 in the control of proliferation and development of CD8^+^ T cell subsets.

Our data reveals that energy metabolism was reprogrammed in CD8^+^ T cells of ATPIF1-KO mice to support the functional change. To elucidate the underlying mechanisms by which the CD8^+^ T cell activities were enhanced in the ATPIF1 deficiency mice, a series of *in vitro* experiments were conducted in isolated CD8^+^ T cells. In the KO mice, the ATP synthase activity was enhanced in ATP production in the glucose-enriched conditions (such as HG and NG) to promote cell proliferation, which reflects release of the ATP synthase activity from suppression. The release protected the cells from apoptosis in the glucose-deficient condition through preservation of the mitochondrial potential (ΔΨm) at the expanse of ATP, a consequence of the elevated hydrolytic activity of the ATP synthase. This reprogramming provided a protection to CD8^+^ T cells in the tumor microenvironment in support of the immune activity. These observations are in line with the catalytic activities of ATPIF1 in other types of cells ^10, 33^. The role of ATPIF1 remains to be established in the control of apoptosis. In the cell culture, ATPIF1 knockdown was reported to reduce tumor growth by promoting apoptosis from impairment of mitochondrial biogenesis and crista remodeling ^15^. In contrast, ATPIF1 inactivation was reported to enhance tumor growth by inhibition of apoptosis in another study ^10^. The exact reason for the different conclusions remains unknown in the two studies. The experiment conditions, such as glucose concentrations and model systems, may hold a promise to answer the question.

The glucose metabolism was reprogrammed in CD8^+^ T cells in the ATPIF1-KO mice. The reprogramming was observed with an elevation in glucose uptake and glycolysis. GLUT1 is the major glucose transporter in T cells for glucose uptake from the microenvironment ^4^. The increased mRNA of GLUT1 led to an active glucose uptake in the ATPIF1-deficient cells. Glycolysis is required for T cell proliferation and production of cytokines including IFN-γ ^20^. In current study, the enhanced glycolysis was identified in the KO cells by the Seahorse assay and the results were confirmed by the data of targeted metabolomics analysis. Glycolysis remains to be established in the differentiation of memory CD8^+^ T cells although OXPHOS is essential for the maintenance of CD8^+^ T_mem_ cells ^34^. Constitutive glycolysis supports the differentiation of CD8^+^ T_eff_ and enhances the antitumor efficacy of CD8^+^ T cells ^35^. OXPHOS was also enhanced in CD8^+^ T cells of the ATPIF1-KO mice in the glucose metabolism. The reprogramming of glucose metabolism in both glycolysis and OXPHOS provides a basis for the activities of CD8^+^ T_eff_/T_mem_ cells in the KO mice.

Mitophagy provides an alternative source of energy to T cells beyond glucose. Mitophagy, a mitochondrion-specific autophagy, is induced by factors including ATP depletion to gain nutrients (fatty acids, amino acids, etc.) from the damaged mitochondria ^36^. Mitophagy was enhanced in the CD8^+^ T cells of ATPIF1-KO mice to form another layer of metabolic reprogramming. The enhanced mitophagy was observed with elevated PINK1, Parkin and Agt5 proteins in the ATPIF1-KO cells under the glucose-limiting (LG) conditions. Mechanism for the autophagy elevation is related to the ATP depletion, which was observed in the KO cells under the glucose-limiting condition due to the enhanced hydrolysis activity of ATP synthase. The depletion promotes mitophagy through activation of AMPK in the T cell response to virus and bacteria ^37^. The mitophagy promotes CD8^+^ T cell survival in the model of influenza virus infection ^38^. Autophagy is required for the antitumor activity of CD8^+^ T cells ^39^. The enhanced mitophagy gives an extra support to the ATPIF1-KO cells in the maintenance of energy homeostasis.

Current study reveals a novel activity of ATPIF1 in T cells. Inactivation of ATPIF1 promotes the tumor immunity in mice by increasing the CD8^+^ T cell activities in several aspects including proliferation, apoptosis resistance, effector, and memory. The functional change is associated with reprogramming in the glucose metabolism and mitophagy in the maintenance of energy hemostasis. The metabolic reprogramming allows the effector and memory CD8^+^ T cells to gain killing activities against the cancer cells in the tumor microenvironment (Fig. 9). The elevation in ATP synthase activity contributes to the metabolic reprogramming for glycolysis, OXPHOS and mitophagy in the glucose-limiting conditions. These data suggest that ATPIF1 protein inhibits the CD8^+^ cell activities in the physiological conditions. ATPIF1 represents a potential molecular target in the induction of CD8^+^ T cell activities for the cancer immunotherapy.

**Figure 9.**
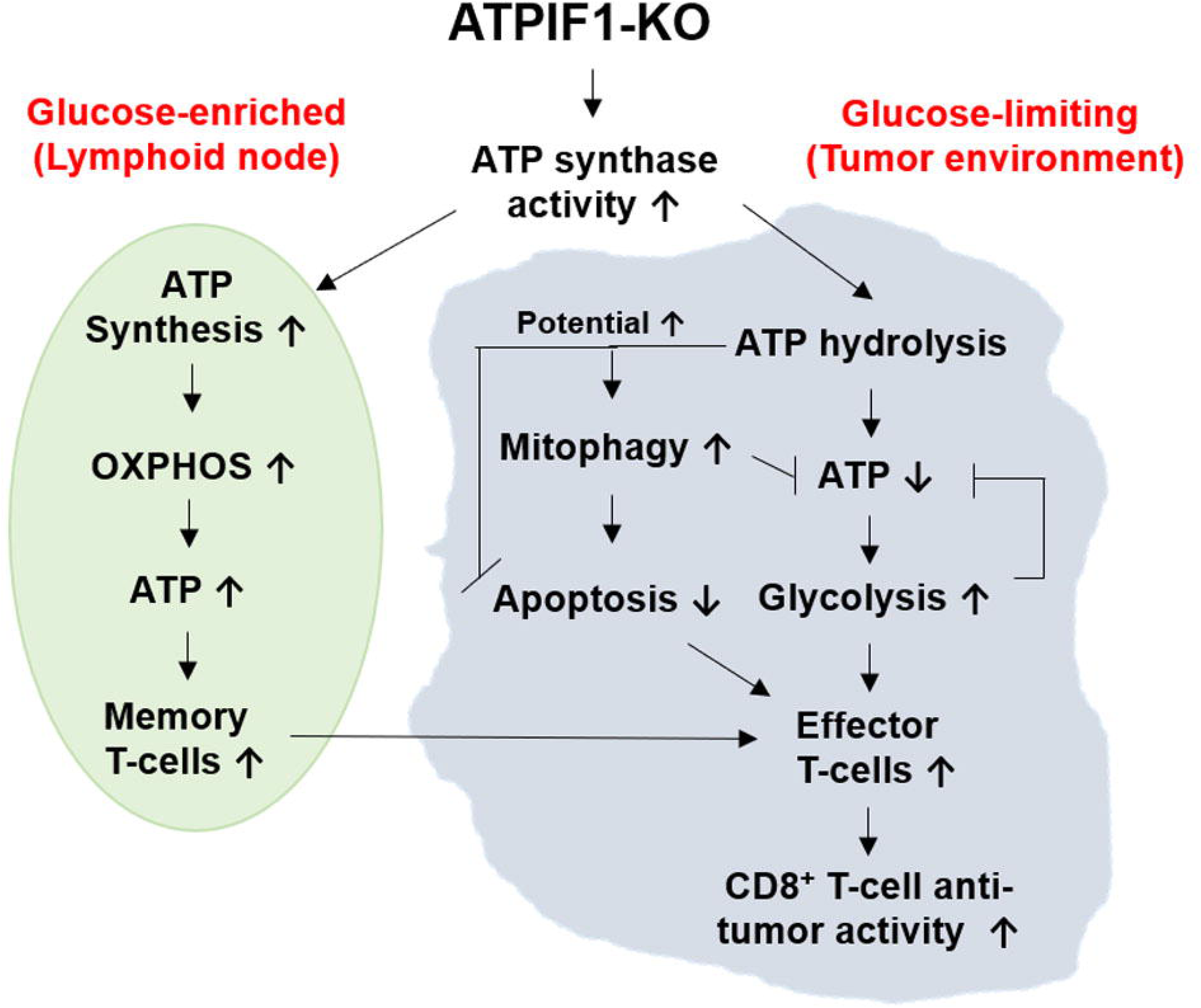
Proposed mechanism for the enhanced antitumor efficacy of CD8^+^ T cells of ATPIF1-KO mice under the glucose-limiting condition.

## Materials and Methods

### ATPIF1^-/-^ mice

The ATPIF1 knockout mouse was generated with the CRISPR/Cas9 gene editing method with sgRNA PAM sequences targeting ATPIF1: SgRNA-1 GCAGTCGGATAGCATGGATACGG and SgRNA-2 GGCTCCACCAGCTTCTCGGATGG. A similar method was described in our published study ^40^. For identification of ATPIF1^-/-^ mice, PCR was conducted to amplify the ATPIF1 gene using the Forward primer CATCAGCCTTGGAATTCTGC and the Reverse primer CTTCGTCTCGGACTCGGTAG. The agarose electrophoresis was performed to determine the genotype, and the amplified PCR product was used in gene sequencing with ABI-3730XL to confirm the genotype. The ATPIF1^-/-^ mice were maintained in the pathogen-free animal facility with free access to food and water, temperature of 20±2 °C, humidity of 60±5%, 12 h light/dark cycle/day at the Sixth People’s Hospital, Shanghai Jiao Tong University, Shanghai, China. All the animal procedures were approved by the Institutional Animal Care and Use Committee (IACUC) of the Shanghai Jiao Tong University and Xinxiang Medical University. Male and female ATPIF1^-/-^ mice at ages of 6-10 weeks were used in this study with the wild type (WT) littermates in the control. The gender-matched WT mice were used in the control.

### Tumor growth and tumor-infiltrating T cells

Mice were implanted subcutaneously with 1×10^6^/mouse of B16-F0 melanoma (CRL-6322™, ATCC), Lewis lung cancer (CRL-1642™, ATCC) or B16-OVA melanoma cells (day 0) in the right axilla to establish the tumor-bearing mouse model. Tumor growth was monitored twice/week, and the tumor size was determined by tissue weight after collection from each mouse at about 2 weeks, when the tumor dimension reached about 20 mm in any direction. In the CD8 T cell deletion experiment, the CD8 antibody was injected intraperitoneally at the dosage of 200 μg/mouse (BE0004-1, BioXcell) at the same time of tumor engraftation. The CD8 antibody was injected twice/week with the control IgG injected in the control group. The tumor-infiltrating lymphocytes were investigated in the excised melanoma tumors (2 mm^3^ tissue) following digestion with the type II gelatinases (0.5%) for 30-40 minutes at 37□ on a shaker. The cell suspension of tumor tissues was filtered with a 70 μm filter obtain the single cell solution. The cells were washed twice with PBS containing 2% FBS and were blocked with CD16/32 Fc block (Cat. 553142, BD Bioscience) before the analysis. The cell populations were examined with different antibodies using the flow cytometry.

For adoptive T cell transfer experiment, the 2×10^6^ B16-OVA melanoma were inoculated into the right flank of Rag1 mice (n=5, purchased from the GemPharmatech Co., Ltd in China), and 9 days after tumor inoculation, 2×10^6^ CD8^+^ OT-I T cells or CD8^+^ ATPIF1^-/-^ OT-I T cells were intratumor injected in 50 μl of saline, respectively. Tumors were measured with a precision caliper every 3 days and the tumor size were calculated.

### Analysis of CD45^+^ tumor-infiltrating lymphocytes by single cell RNA-seq

B16 melanoma cells were implanted in the right flank of male WT and KO mice (n=6). On day 12, the tumors were dissected from the surrounding fascia, mechanically minced to approximately 1 mm^3^ pieces, then the tumor tissue suspension from KO mice (n=6) was mixed together, so did the tumor tissue suspension from WT mice. After that, the mixed tumor tissue suspension was stored in a solution provided by the Beijing Analytical Biosciences Technology Co., Ltd, a company specialize at single cell RNA sequencing and data analysis, which is founded by the Professor Zhang Zemin in Peking University, Beijing, China. The samples of WT and KO mice were sent to this company for CD45^+^ T cell sorting immediately and single cell RNA-seq. The preparation of tumor single cell suspension, the CD45^+^ tumor-infiltrating leukocytes sorting, reverse transcription, amplification, sequencing, the quality control, scRNA-seq data processing, cluster analysis, trajectory analysis, STARTRAC analysis and RNA velocity-based cell fate tracing were all performed in the Beijing Analytical Biosciences Technology Co., Ltd, which was described in detail in two studies ^16, 41^.

### Glucose uptake

The CD3^+^ T cells were collected and cultured in the high glucose (HG, 4.5 g/L glucose), low glucose (LG, 0.2 g/L glucose) and normal glucose (NG, 2 g/L glucose) conditions for 24 h, then resuspend in glucose-free medium with the addition of 2-NBDG (20 μM). After incubation for 1 h at 37□, the T cells were stained with CD3 and CD8 antibodies, and propidium iodide (PI). Glucose uptake was determined with 2-NBDG quantification using the flow cytometry. The ratio of 2-NBDG positive and negative CD8^+^ T cells was calculated and presented.

### Mitochondrial function and glycolysis

CD8^+^ T cells were isolated from the mouse spleen and activated with CD3/CD28 antibody stimulation in the culture medium. The cells were loaded at 1 × 10^6^/well into XF24 plate, which was coated with Cell-Tak (company name, 22.4 μg/mL, in sterile water) for 20 minutes to increase the adhesion of T cells. The plate was centrifuged at 200 *g (zero braking) for 1 minute to let the T cells adhere to the culture surface. After incubation for 30 minutes at 37 □without CO_2_ supplementation, Oxygen Consumption Rate (OCR) and Extracellular Acidification Rate (ECAR) were determined with the Seahorse XF Cell Mito Stress Test Kit and Seahorse XF Glycolysis Stress Test kit with the Agilent Technologies equipment. The final concentrations of inhibitors were: 1 μM oligomycin, 2 μM FCCP, 0.5 μM rotenone and antimycin A. In ECAR assay, the final concentrations of compounds were: 10 mM glucose, 1 μM oligomycin, and 50 mM 2-Deoxy-D-glucose. The readings were taken after each sequential injection of corresponding chemicals.

### Targeted metabolomics

The CD8^+^ T cells were isolated from the spleen of tumor-bearing mice, which were inoculated with B16-OVA tumor for 2 weeks (n = 6), and frozen immediately with dry ice. The energy metabolites were quantified using the LC-MS method. The analysis included 32 major metabolites of the tricarboxylic acid cycle (TCA), glycolytic pathway, pentose phosphate pathway and oxidative phosphorylation pathway. The hierarchical clustering and quantity of the metabolites are presented to show the change in lymphocytes.

### MitoTracker assay

T cells were cultured in the high glucose (HG, 4.5g/L) medium, the low glucose (LG, 0.2g/L) medium and normal glucose (NG, 2g/L) medium. The mediums were prepared in glucose-free RMPI-1640 medium supplemented with glucose, 10% FBS, 1% Penicillin-Streptomycin solution. After culture for 24 h, the mitochondrial mass MitoTracker (Cat. M7514, Thermo Fisher Scientific) were determined according to the manufacturer’s protocol with flow cytometry (BD FACS Calibur).

### Mitochondrial membrane potential (ΔΨm)

The T cell were cultured under high glucose (HG, 4.5 g/L glucose), low glucose (LG, 0.2 g/L glucose) and normal glucose (NG, 2 g/L glucose) in the RMPI-1640 medium with the addition of JC-1 in the Mitochondrial membrane potential assay kit (Beyotime Biotechnology Limited Company. Shanghai, China), and incubated in the normal (O_2_ concentration about 20%) or the low oxygen (1%) condition. After 24 h culture, the T cells were collected and stained with CD3-APC/CD8-PerCP antibodies and analyzed with flow cytometry.

### Western blotting

To detect the protein expression, the CD8^+^ T cells were cultured in the HG, NG and LG condition and lysed in RIPA with the proteinase inhibitor cocktail. The Western blotting was conducted according to a published protocol ^42^. The primary antibodies to Glut-1 (#12939, CST), PINK1(Cat. Ab186303, Abcam), Parkin (Cat. Ab77924, Abcam), VDAC1 (ab154856, Abcam), ATG5 (#12994, CST) and β-actin (Cat. AC026, ABclonal) were used to blot the membrane for corresponding proteins.

### T cell proliferation and mitophagy

CD3^+^ T cells were isolated from the spleen of C57BL/6 mice using a negative selection T cell separation kit (#8802-6840-74, Thermo Fisher Science). CD3^+^ T cells were labeled with the CellTrace carboxyfluorescein diacetate succinimide ester (CFSE; ThermoFisher, Cat. C34554), and then cultured with 30 U/mL rIL-2 (#34-8021-85, eBioscience) and activated with the mouse T-activator CD3/CD28 Dynabeads (Cat. 11452D, ThermoFisher) for 96 hours under the HG, NG, and LG conditions. The T cells were collected and stained with CD3-PE, CD8-PerCP and then analyzed with the BD Calibur FACS for evaluation of cell proliferation. For mitophagy activation experiment, the CD8^+^ T cells were cultured under the LG condition for overnight, then treated with CCCP (100 μM, #B5003 APExBIO) for additional 3 h under in the LG condition. The pellets were collected for Western blotting (PINK1, Parkin, VDAC1 and β-actin) or analysis of IFN-γ with FACS analysis following a fix/permeabilization procedure.

### Cytokine assay

The concentration of IFN-γ was determined in the cultural medium with an ELISA kit according to the manufacturer’s protocol (Cat. 88-7314-22, ThermoFisher). In CD8^+^ T cells, the flow cytometry was used to detect INF-γ in the CD3^+^ T cells after activation with the eBioscience™ Cell Stimulation Cocktail plus protein transport inhibitors (Cat. 00-4975-93, 500X). The cells were stained with CD3-PE and CD8-FITC antibody, then fixed with the eBioscience™ Fixation/Permeabilization Diluent agents (Cat. 00-5223-56) for 1 h. After washing out the two antibodies, the cells were stained with IFN-γ-APC antibody and then analyzed with the FACS station.

### qRT-PCR

mRNA was extracted from the CD8+ T cells using the TRIzol agents (TAKARA, Japan). mRNA expression was quantified in qRT-PCR using the TB Green® Premix Ex Taq™ II (Tli RNaseH Plus) for cDNA generation, and the PrimeScript™ RT Master Mix (Perfect Real Time) in PCR analysis. The primers include: (Slc2a1-Forward: 5’-TGGGCAAGTCCTTTGAGATG-3, Slc2a1-Reverse: 5’-TGACACCTCTCCCACATACA-3; pdk1-Forward: 5’-GGGACAGATGCGGTTATCTAC-3, pdk1-Reverse: 5’-CTCGTGGTTGGCTTTGTAATG-3; β-actin-Forward: 5’-CCACACCCGCCACCAGTTCG-3, β-actin-Reverse: 5’-TACAGCCCGGGGAGCATCGT-3). The qRT-PCR reaction was conducted using the 7500 Fast Real-Time PCR station (Applied Biosystems). The β-actin mRNA was used as internal reference in data normalization. The results are presented in mean ± SD of 3-6 samples.

### Coculture of OT-I CD8^+^ T cells with B16-OVA tumor

The ATPIF1^-/-^ OT-I CD8^+^ T cell and OT-I CD8^+^ T cells were isolated from the spleen of ATPIF1^-/-^ OT-I and OT-I mice, and cocultured with B16-OVA cell under high glucose (4.5 g/L glucose), normal glucose (2 g/L glucose), low glucose (0.2 g/L glucose) in 10% FBS contained RMPI-1640 medium in regular O_2_ concentration (∼20% O_2_) incubator for 36 h, then the IFN-γ secretion of CD8^+^ T cell and CD8^+^ T_CM_ were determined. For IFN-γ determination, the coculture CD8^+^ T cells were collected and labeled with CD3-FITC, CD8-PE, after PBS washed for one time, the pellets were fixed and permeabilized using the FIX & PERM™ Cell Permeabilization Kit purchased from Thermofisher (Cat. GAS004). Then the IFN-γ-APC was stained for detection of intracellular IFN-γ expression. For T_CM_ detection, the CD3, CD8, CD44 and CD62L antibody were used to stain the collected T cells, then the CD62L^+^CD44^+^ CD8^+^ T cells were indicated as T_CM_ cells and CD62L^-^CD44^+^ CD8^+^ T cells were indicated as T_EM_ cells. In the coculture process, the apoptosis was also analyzed using the Annexin V and 7-AAD. Firstly, the collected T cells were labeled with CD3-APC, CD8-FITC, Annexin V-PE and 7-AAD for detection of apoptosis according to the protocol, then loaded onto the BD Calibur FACS for analysis.

## Supporting information

Fug S6

## Data availability

All the raw data of scRNA-seq and the TCR sequencing were uploaded to the GEO with the accession number of GSE158278. (https://www.ncbi.nlm.nih.gov/geo/query/acc.cgi?acc=GSE158278)

## Statistical analysis

The quantitative data are presented as the mean ± SD. Unpaired two-tailed t-test was used in the data analysis for a difference between the WT and KO groups, and *p* value < 0.05 was considered statistically significant. FlowJo 10.0 was used to analyze the flow cytometry results.

## Acknowledgements

This work was supported in part by the Natural Science Foundation of China (No. U1904131, No. 81872361) and the Key Scientific and Technological Research Projects in Henan Province (No. 192102310169)

## Author contributions

G.Z. and J.Y. designed the study and wrote the manuscript. G.Z., Y.W., J.Z., Y.W., Y.G., S.S., X.Z., X.C., Y.L., M.W., Z.Z., M.S. and Y.L. conducted the experiments and data analysis. Y.L. and H.W. generated the ATPIF1 KO mice and provided technical support in verification of ATPIF1-KO mice.

## Competing interests

All the authors declare no competing interests.

## Figure legends

**Figure S1**. Lymphocyte population analysis in WT and ATPIF1^-/-^ mice. (a) CD4 and CD8 populations in thymus. (b) Lymphocyte populations of spleen. CD5^+^ T-cells, CD4^+^ T cells, CD8^+^ T cells, B cells, Tregs, dendritic cells and CD44^+^CD62L^-^ memory T cells were determined in spleen. (c) Bar figures of lymphocyte populations in the spleen. * *p*<0.05 (n=5).

**Figure S2**. Representative graph of detecting CD8 T cell. The CD8 T cells were deleted in mice via intraperitoneal injection of CD8 antibody at 200 μg per mice in three days. (a) WT mice without CD8 antibody injection. The blood was analyzed using the flow cytometry. (b) WT mice with CD8 antibody administration. Data indicated the CD8 T cells were effectively deleted after the administration of CD8 antibody.

**Figure S3**. Sorting of CD45^+^ leucocytes from the tumor single cell suspension. The tumors excised from the WT and KO mice were digested to obtain the single cell suspension and then subject to sorting by the flow cytometry. The BV421 was used to sort the live and dead cells, BV421 negative cells was gated for the CD45^+^ sorting as indicated in WT and KO sample.

**Figure S4**. Quality control of the scRNA-seq indicated by the violin chart (a), PCA (b), tsne and UMAP (d) analysis.

**Figure S5**. Heatmap of the marker genes in eight clusters of T cell subsets based on 10x genomics sequencing.

**Figure S6**. Heatmap of the marker genes in four major different clusters of B cells, DCs, Macrophages and T cell subsets based on 10x genomics sequencing.

**Figure S7**. The pseudotime trajectory analysis of DC subsets based on the scRNA-seq results. (a) The order of DC cells along pseudotime in a two-dimension state-space defined by Monocle2. Cell orders were inferred from the expression of most dispersed genes across DC subsets sans MAIT. Each point indicates a single cell, and each color represents a DC cell cluster. (b) The pseudotime trajectory of DC populations in WT and KO samples. The red dot is the KO sample and green dot is the WT sample.

## Tables

**Table S1**. Sample statistics after quality control during the 10×genomics single cell sequencing.

**Table S2**. Summarized data in Excel format of the targeted metabolomics of the WT and KO CD8^+^ T cells.

## Notes

https://www.ncbi.nlm.nih.gov/geo/query/acc.cgi?acc=GSE158278

### Competing Interest Statement

The authors have declared no competing interest.

