## Supplementary material for "ATPIF1 inactivation promotes antitumor immunity through metabolic reprogramming of CD8^+^ T cells": Table 1

| **Table S1. Sample information** | | |
| --- | --- | --- |
|  | WT | KO |
| Cell count raw | 10059 | 8408 |
| Cell count filtered | 8010 | 6007 |
| UMI counts raw | 994 | 654 |
| UMI counts filtered | 1422.5 | 1324.0 |
| Gene counts raw | 467 | 339 |
| Gene counts filtered | 617 | 614 |
| Mito percent raw | 2.22 | 3.95 |
| Mito percent filtered | 2.1 | 2.6 |
