## Supplementary figures and images for "ATPIF1 inactivation promotes antitumor immunity through metabolic reprogramming of CD8^+^ T cells"

### Fig S1

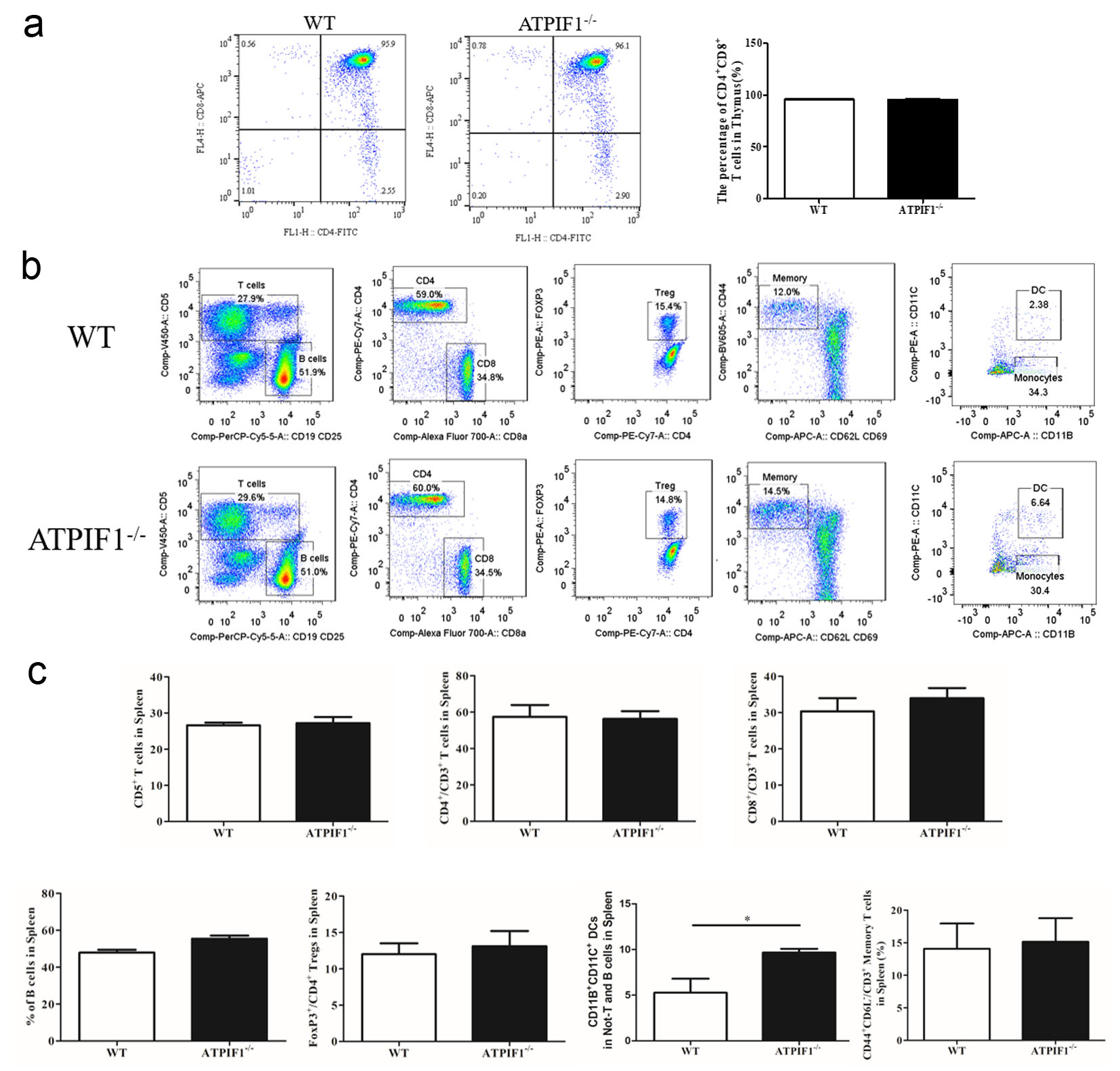

### Fig S2

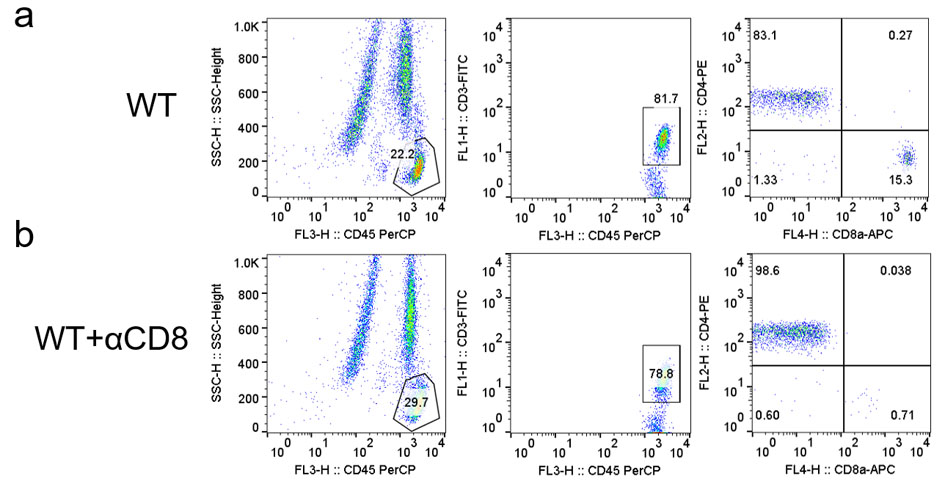

### Fig S3

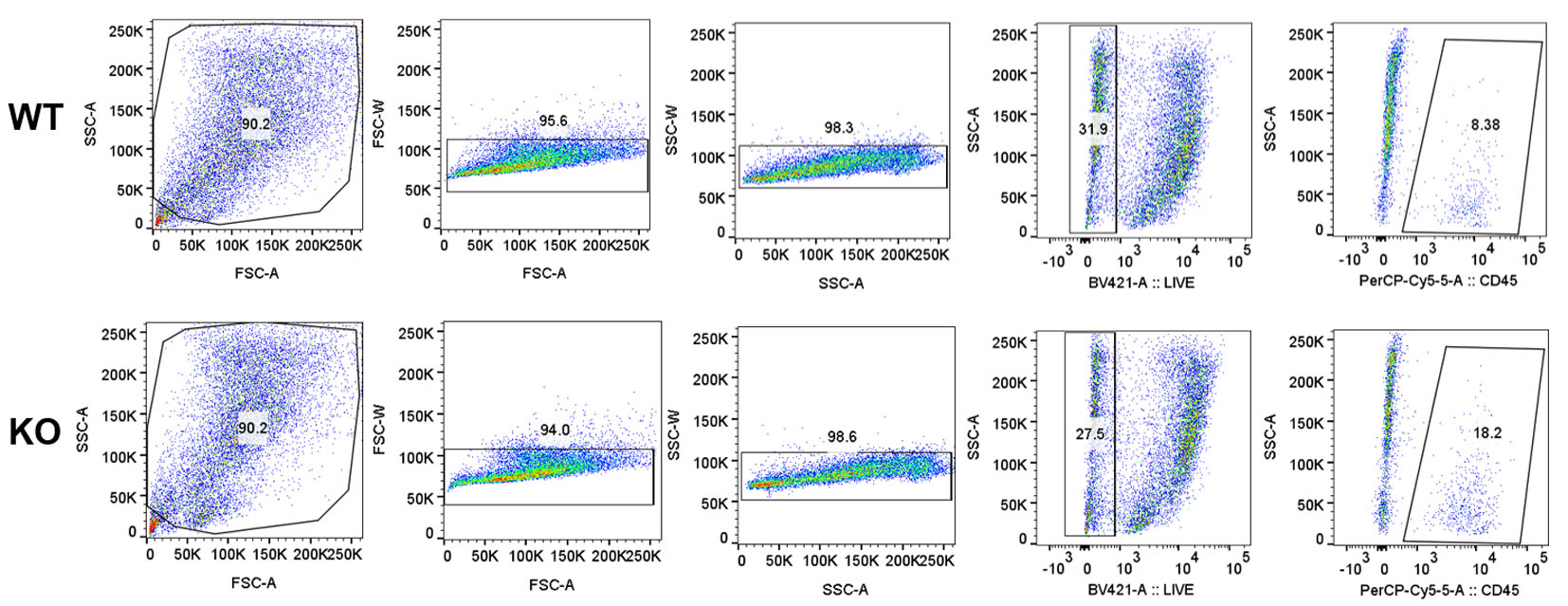

### Fig S4

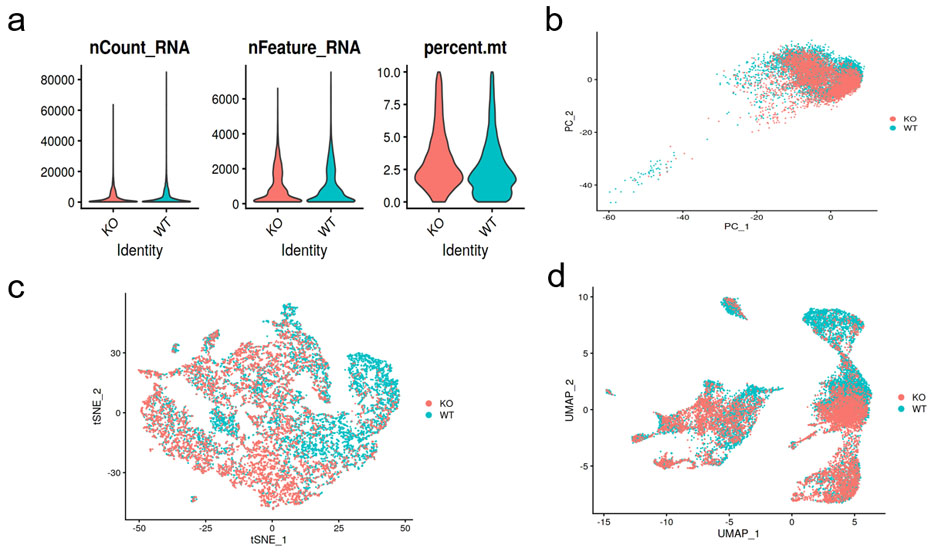

### Fig S5

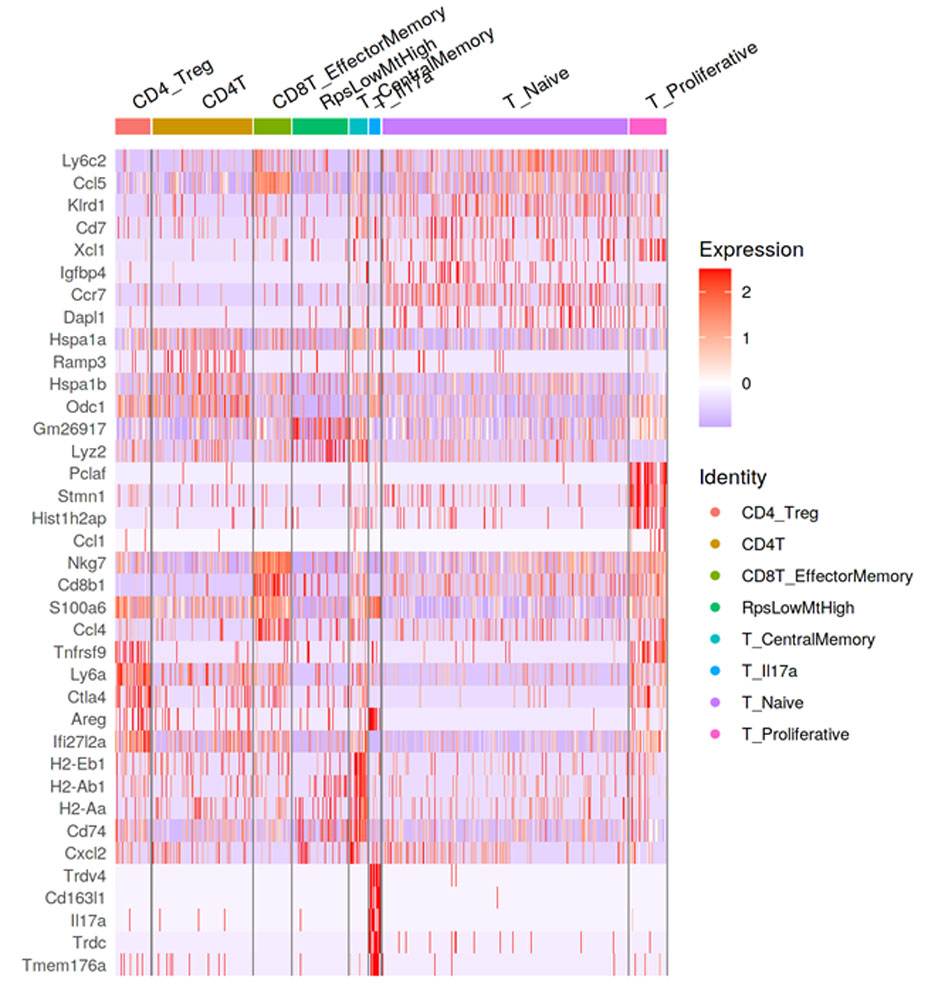

### Fig S7

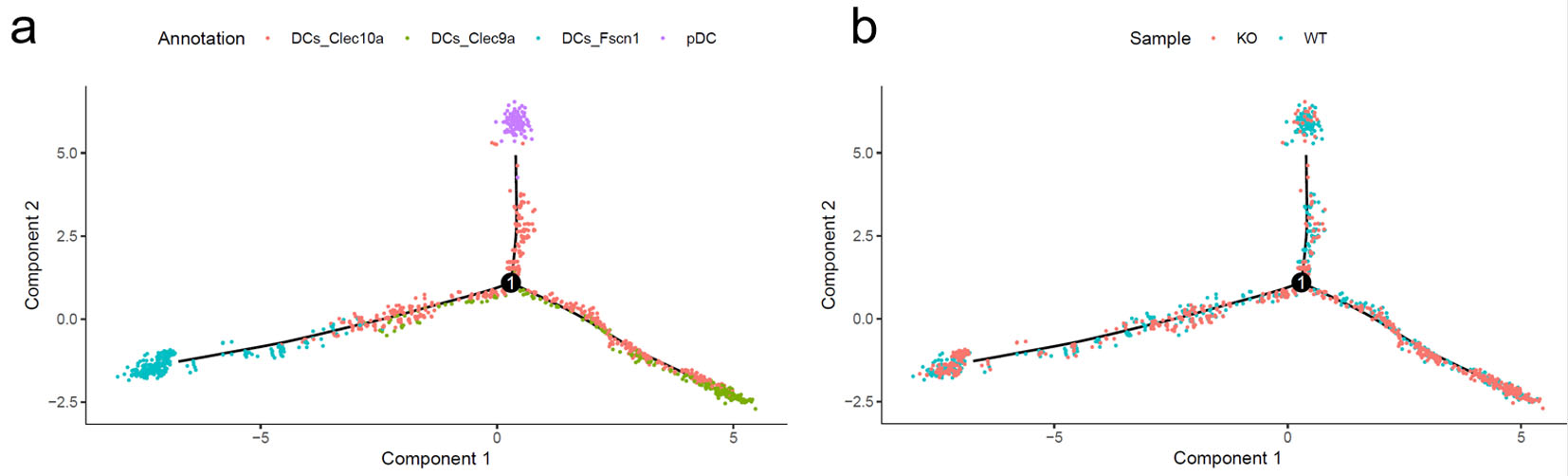

### Fug S6

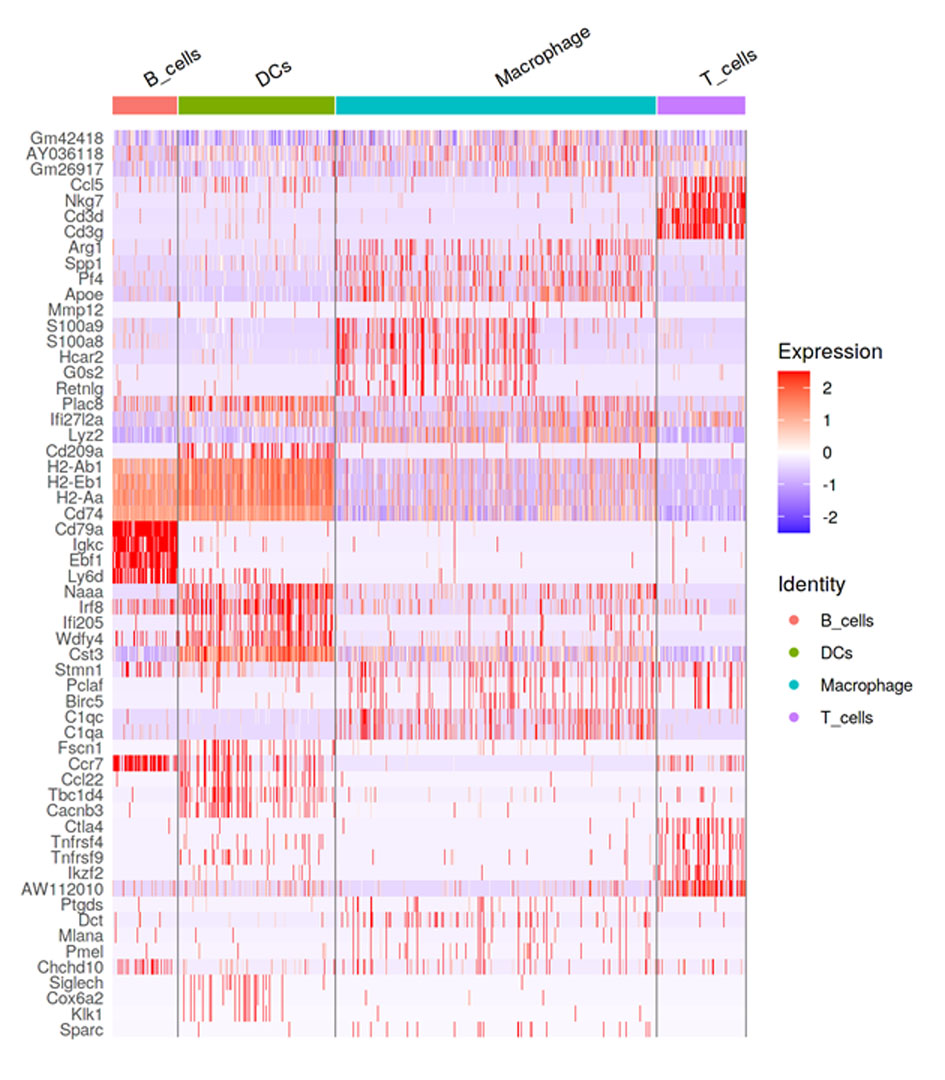
